# SCGN Administration prevents Insulin Resistance and Diabetic Complications in High-Fat Diet Fed Animals

**DOI:** 10.1101/189324

**Authors:** Anand Kumar Sharma, Radhika Khandelwal, Swathi Chadalawada, N Sai Ram, T Avinash Raj, M Jerald Mahesh Kumar, Yogendra Sharma

## Abstract

Secretagogin (SCGN) is poorly-studied secretory/cytosolic CaBP enriched in pancreatic **β**-cells. Recent studies implicated SCGN in diabetes; however, its function and therapeutic prospect remain uncharted. Based on the apparent synchrony of SCGN and insulin secretion (and its disruption in HFD-fed animals) and considering *SCGN* downregulation in Type 2 diabetes, we hypothesized that SCGN is a key regulator of insulin response. To test this, we administered rSCGN to HFD-fed animals. We here report that a novel SCGN-insulin interaction stabilizes insulin and potentiates insulin action. Moreover, a chronic rSCGN administration improves insulin response and alleviates obesity associated risk factors such as weight gain, liver steatosis and cholesterol imbalance in DIO animals. Beside the anti-diabetic effects, prolonged rSCGN treatment also induces β-cell regeneration. These effects seem to originate from SCGN mediated regulation of insulin concentration & function as validated in insulin-deficient STZ animals. Our results demonstrate the prospects of the therapeutic potential of SCGN against diabetes.

## Introduction

With the distressing growth in the diabetes cases worldwide, the search for endogenous therapeutic candidates has become a prevailing biomedical research theme (1-5). The risk of drug resistance, associated side-effects of currently available therapies and prevalence of genetic unresponsiveness to drugs further emphasizes the need to explore alternate candidates. In this line, we noticed Secretagogin (SCGN; a six-EF hand Ca^2+^-binding protein) as an intriguing prospect. This is based on the facts that in pancreatic β-cells, SCGN regulates both the insulin expression (6) and the release (6-10). Moreover, SCGN seems to be synchronously secreted with insulin upon cAMP stimulation (11). Likewise, a measurable fraction of SCGN dwells in the circulation (12-13) but the biological functions of circulatory SCGN remain unknown.

Based on the available information and its logical acquaintance with SCGN function, we predicted the involvement of SCGN in the regulation of obesity and insulin resistance. Association of *SCGN* with body-weight QTC (14), downregulation of SCGN in HFD fed animals (7) and reduced CSF SCGN concentration in the insulin resistant subjects (15) offer confidence to our rationalized assumption. The development of glucose intolerance at an early life-stage and age associated progressive hyperglycemia in SCGN knockout mice further suggest a confounding role of SCGN in preserving euglycemia (16). Moreover, reduced SCGN expression in T2DM patients and a negative correlation of SCGN expression with HbA1c (16) suggest a poised but perplexing association of SCGN with diabetes (particularly insulin resistance). A broader and applied interest is to scrutinize SCGN’s therapeutic potential against insulin resistance. Considering these observations, we hypothesized that because SCGN deficiency correlates with the incidence of diabetes, the exogenous rSCGN administration will have an anti-diabetic effect *in vivo*.

We demonstrate here that exogenous SCGN can be used as a therapeutic agent in diabetes. In this maiden study on the extracellular function of SCGN, we report our observation of extracellular SCGN-insulin interaction and it's biological implication in increasing the potency of insulin *in vivo* which then prompted us to explore therapeutic use of SCGN as an insulin sensitizer/fortifier. We found that a chronic SCGN administration has insulin sensitizing activity in DIO animals. Our results designate SCGN as an anti-diabetic protein. SCGN’s anti-diabetic activity plausibly originates both from insulin potentiation as well as tissue remodeling to increase insulin sensitivity (of insulin target tissues) and optimal insulin synthesis and secretion. Relating previously known literature and our current findings, it is convincing clear that SCGN not only regulates insulin release but also its expression, stability and functions beyond pancreas. We suggest that recombinant SCGN (or SCGN derived smaller active peptide) emerges as a potential anti-diabetic monotherapy and can also be used as combination therapy.

## Results

### SCGN readily interacts with insulin: the synergy of Ca^2+^ and insulin binding

Apart from the role of SCGN in insulin, other pancreatic β-cell specific functions and the circulatory functions of SCGN remain undiscovered. To explore for novel SCGN interaction and function, we performed a mass-spectroscopy based pull-down assay from MIN6 cell lysate. We identified several Ca^2+^ binding/chaperone proteins (which are implicated in insulin maturation process) as SCGN interactors (e.g., BiP, PDI, GRP94) (Fig. 1a). Interestingly, in addition to the aforesaid chaperones, we observed that insulin was also captured as a SCGN-interactor. Considering previously reported occasional colocalization of SCGN with insulin (6, 11), we checked for the targeted colocalization in a stable SCGN-eGFP expressing MIN6 cell line. We observed that SCGN (green) and insulin (red) colocalize (yellow) to a substantial magnitude (Fig. 1b). To validate SCGN-insulin interaction in the extracellular milieu, we performed a protein overlay assay and observed that SCGN binds insulin, irrespective of the presence of Ca2+ (Fig 1c). Even partially unfolded insulin (insulin-treated with 4 M urea prior to immobilization) binds to SCGN suggesting a possibility of SCGN acting as a chaperone for insulin. We also performed a CoIP assay from MIN6 cell lysate to confirm the in-cell interaction. We observed a co-precipitation of SCGN with insulin (Fig. 1d).

**Figure 1.**
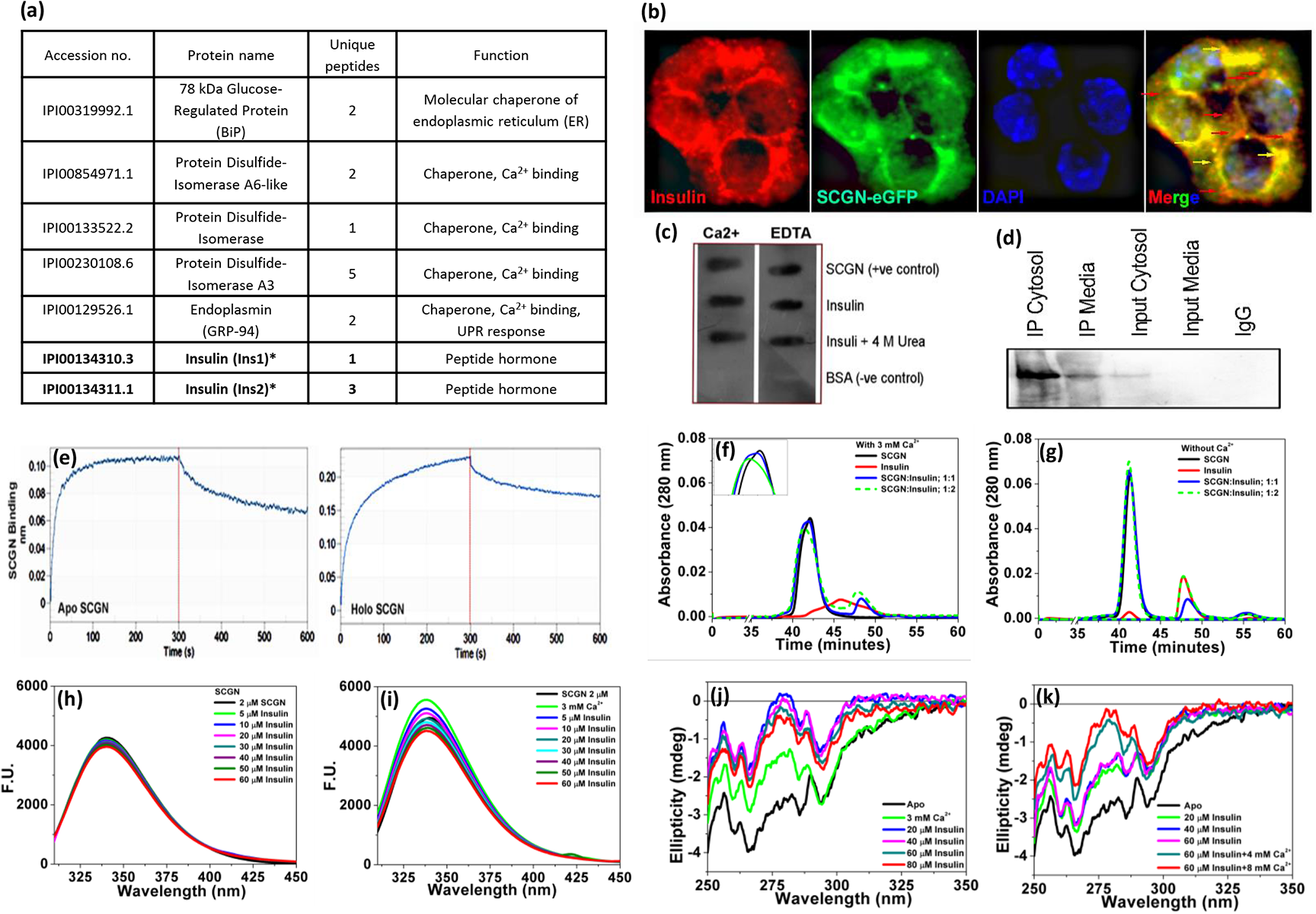
SCGN interacts with insulin *in vitro* and *ex vivo*. (a) List of chaperones (and insulin) identified as SCGN interacting partners in mass spectrometry assisted pull-down assay; (b) Microscopy images depicting colocalization (yellow) of eGFP-SCGN (green) and insulin immuno-stained with Alexa Fluor 594 (red) in MIN6 cells; (c) Protein overlay assay to assess the SCGN-insulin interaction; (d) CoIP from MIN6 cell lysate - anti-insulin immunoprecipitation; (e) BioLayer Interferometry (BLI) binding curve of SCGN-insulin interaction. The data were fit in 1:1 model. R^2^ for apo-SCGN-insulin experiment is 0.9547 (Chi^2^ value is 0.004) and for holo-SCGN-insulin binding is 0.9626 (Chi^2^ value being 0.0126); (f,g) analytical size exclusion chromatography of SCGN alone, insulin alone and SCGN-insulin mixture in 1:1 and 1:2 stoichiometric ratios in the absence and presence of Ca^2+^; (h,i) fluorescence spectra demonstrating changes in Trp microenvironment of SCGN upon insulin titration into the apo- and holo-proteins; (j,k) Near UV-CD spectra of apo- and holo-SCGN upon insulin binding illustrate global conformational changes in the protein.

Being a Ca2+ sensor, a large degree of functional regulations of SCGN emanates from Ca2+ binding. To evaluate the effect of Ca2+ on insulin binding, we performed bio-layer interferometry (BLI) analysis (Fig. 1e). Apo-SCGN (Ca2+-free SCGN) binds insulin with an affinity in the lower micromolar range (Kd=1.84 µM) while the presence of Ca^2+^ slightly increases the overall binding affinity (Kd=1.45 µM). The rate of SCGN-insulin complex formation (k*on*) is of the same order irrespective of the presence of Ca^2+^ (k_on_ for apo-SCGN= 7.21x10^2^; for holo-SCGN= 4.64x10^2^). On the contrary, the rate of dissociation in the presence of Ca^2+^ is decreased by an order of magnitude (k_off_ for apo-SCGN=1.33x10^-3^, for holo-SCGN=6.74x10^-4^) suggesting a strong and specific binding. In the biological context where cytosol has nM [Ca^2+^] while the extracellular milieu has mM [Ca^2+^], the apparent lack of Ca^2+^ dependency would make the interaction pervasive.

We further analyzed the solution behavior of the insulin-SCGN complex. In analytical gel filtration, the holo-SCGN-insulin mixture (but not apo-SCGN-insulin) elutes prior to the holo-SCGN peak suggesting the formation of a ternary complex between Ca^2+^-SCGN and insulin (Fig 1f, g). Similarly, the titration of insulin to holo-SCGN results in a more prominent reduction in the fluorescence intensity than apo-SCGN suggesting that insulin binding is stronger when SCGN is complexed with Ca^2+^. The addition of Ca^2+^ to apo-SCGN-insulin complex results in an increase in fluorescence intensity with a distinct blue shift. However, when insulin is added to the holo-SCGN, insulin binding does not oppose the blue shift (Fig 1h, i). Since the blue-shifted spectrum is a hallmark of Ca^2+^-bound SCGN (17-18), the apparent absence of λ max shift by insulin to holo-SCGN suggests that Ca^2+^ is neither replaced nor removed allosterically by insulin. These results also suggest that the insulin and Ca^2+^ binding sites are mutually independent.

Mutual independence of Ca^2+^ and insulin binding is further supported by near-UV circular dichroism spectral analysis. Ca^2+^ binding induces a large conformational change in the aromatic region of SCGN tertiary fingerprint (17-18). Remarkably, insulin binding to apo-SCGN induces similar changes in near-UV CD spectra (between 260-290 nm) (Fig. 1j, k). Upon addition of Ca^2+^ to apo-SCGN-insulin complex, a sharp decrease in the negative ellipticity (around 280 nm) takes places, while the identity of all the major aromatic CD bands is preserved, demonstrating a drastic alteration in the tertiary conformation of SCGN-insulin complex by Ca^2+^ (Fig. 2e, f). Incidentally, insulin induces similar changes in near-UV CD spectrum of Ca^2+^-SCGN producing similar conformational changes irrespective of the order of the ligand.

**Figure 2.**
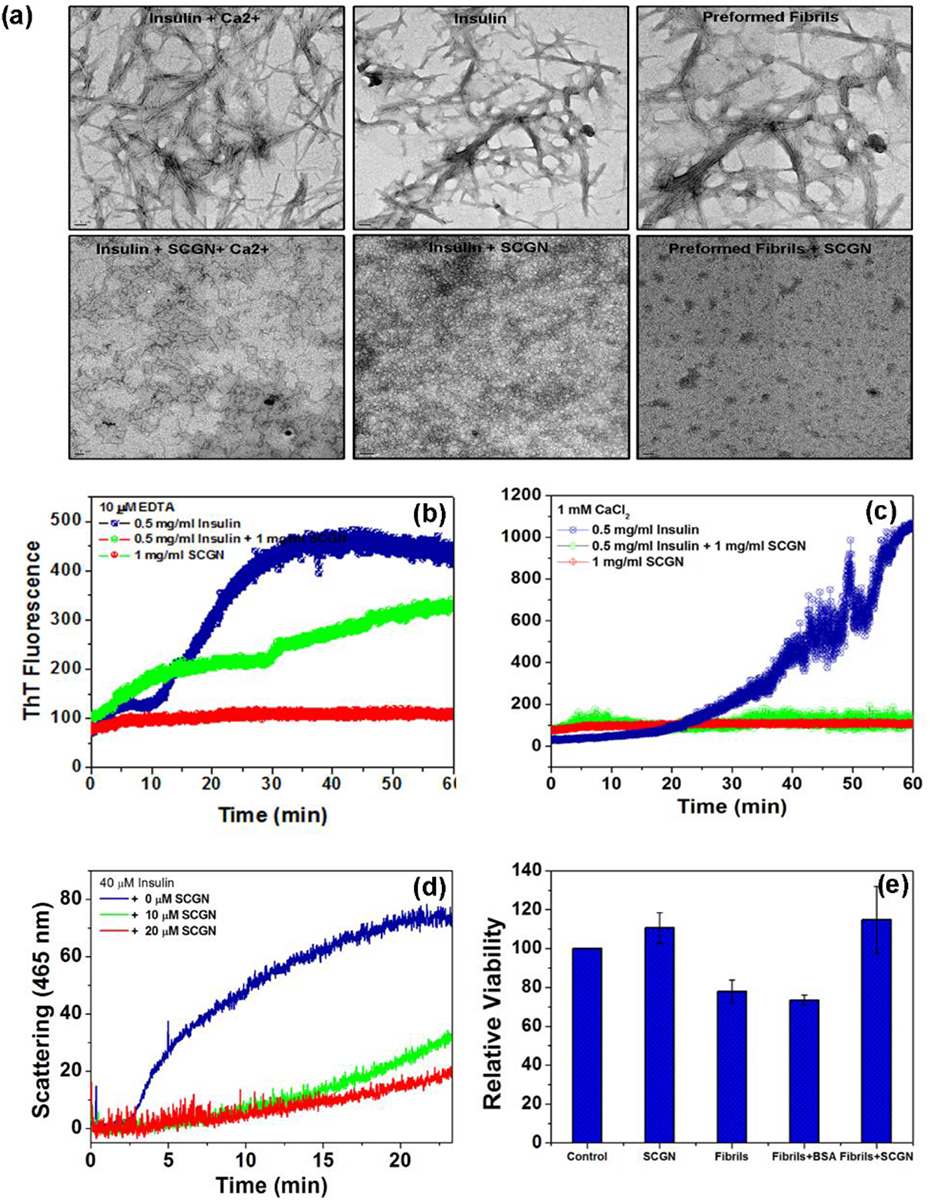
SCGN protects insulin from aggregation and fibrillation and reduces cytotoxicity *ex-vivo*. (a) Transmission Electron Microscope (TEM) images depicting the inhibition of insulin fibrillation by SCGN; (b-c) Time scan spectra representing the change in the emission of ThT fluorescence of insulin fibrillation in the presence of SCGN (1:2 ratio) in its apo- and holo-forms; (d) Change in Rayleigh scattering demonstrating the extent of insulin aggregation upon addition of DTT; (e) MTT cell viability assay demonstrating the role of SCGN in alleviating cell cytotoxicity caused due to insulin fibrils in MIN6 cells. (fibrils vs fibril+SCGN p value= 0.034).

### SCGN protects insulin from aggregation and fibrillation and alleviates fibril toxicity

The major regulation of insulin action is supposedly achieved at the insulin secretion level. The structural stabilization and activity modulation of circulatory insulin, however, remain poorly understood. Since insulin has a soaring propensity to form cytotoxic fibrils, we speculated that SCGN would protect insulin from such chaos. Consistently, we observed that SCGN impedes heat-induced fibrillation of insulin (Fig. 2). The inhibition of the nucleation of fibrillation is likely independent of Ca^2+^, as the apo- as well as holo-SCGN, inhibits the insulin fibrillation (Fig. 2a; vertical panels 1 and 2). SCGN could also dissolve the preformed fibrils within an hour under given experimental conditions (Fig. 2a; last panel). We then measured the rate of fibril formation by ThT fluorescence. We observed that apo-SCGN exerts ~50% protection against heat-induced amyloidogenesis of insulin (Fig. 2b), while holo-SCGN completely abolishes the fibrillation of insulin (Fig. 2c). The protection exerted by SCGN is also visible in the disulfide-bond reduction induced aggregation of insulin. Apo-SCGN rescues insulin from aggregation in a dose-dependent manner (Fig. 2d), implying that apo-SCGN stabilizes insulin, likely by providing a niche to protect the exposed disulfide bonds involved in the interchain disulfide bridge formation. Due to the anomalous behavior of insulin aggregation profile in the presence of Ca^2+^, this experiment could not be performed. Nonetheless, it is likely that Ca^2+^-bound SCGN would be even more effective. These results suggest that SCGN prevents insulin misfolding & aggregation, and chops the existing fibrils. Because of their cytotoxicity, fibrils have been implicated in various neurodegenerative diseases and diabetes (19-24). we wondered if SCGN could alleviate the cytotoxicity caused due to insulin fibrils *ex vivo*. We found that for MIN6 cells, extracellular SCGN reduces insulin fibril toxicity significantly (Fig. 2e). BSA, which was included as a negative control, could not exert any protection. These results suggest that SCGN effectively inhibits the fibrillation of insulin, dissolve the preformed fibrils and alleviate the fibril-associated cytotoxicity. Chaperone interactome and anti-amyloid property of SCGN suggest that SCGN is a molecular chaperone.

### SCGN improves insulin signaling

We next proceeded to study the impact of SCGN on insulin signaling. We found that SCGN enhances the insulin-dependent insulin receptor signaling as evident from increased Akt phosphorylation in two different cell lines (HepG2, C2C12 myotubes) (Fig. 3a). The increased Akt phosphorylation in SCGN-insulin complex treated cells was largely insulin-mediated; however, SCGN alone could also induce slight phosphorylation. Moreover, since internalized insulin (as insulin-IR complex) can activate different pathways to potentiate the receptor mediated effect (25-26), we thus measured the FITC-insulin internalization in the insulin-responsive cell line, HepG2. As seen in the microscopy images, SCGN increases the internalization of FITC-insulin to a visually appreciable level (Fig. 3b). In addition, in the presence of SCGN, there were more visible puncta or granular structures pointing towards improved vesicular endocytosis of insulin-IR complex by SCGN. We validated microscopic data with FACS-assisted quantification of FITC-insulin internalization (Fig. 3c, d). The intensity of the internalized FITC-insulin was higher in the presence of SCGN than in the absence thereof. The results suggest that SCGN possibly facilitates the receptor-binding and might have a role in the hepatic clearance of circulatory insulin. Because a sustained increase in circulatory insulin is a direct correlate of insulin resistance and diabetes, SCGN could impede the progression of insulin resistance by inducing the insulin internalization.

**Figure 3.**
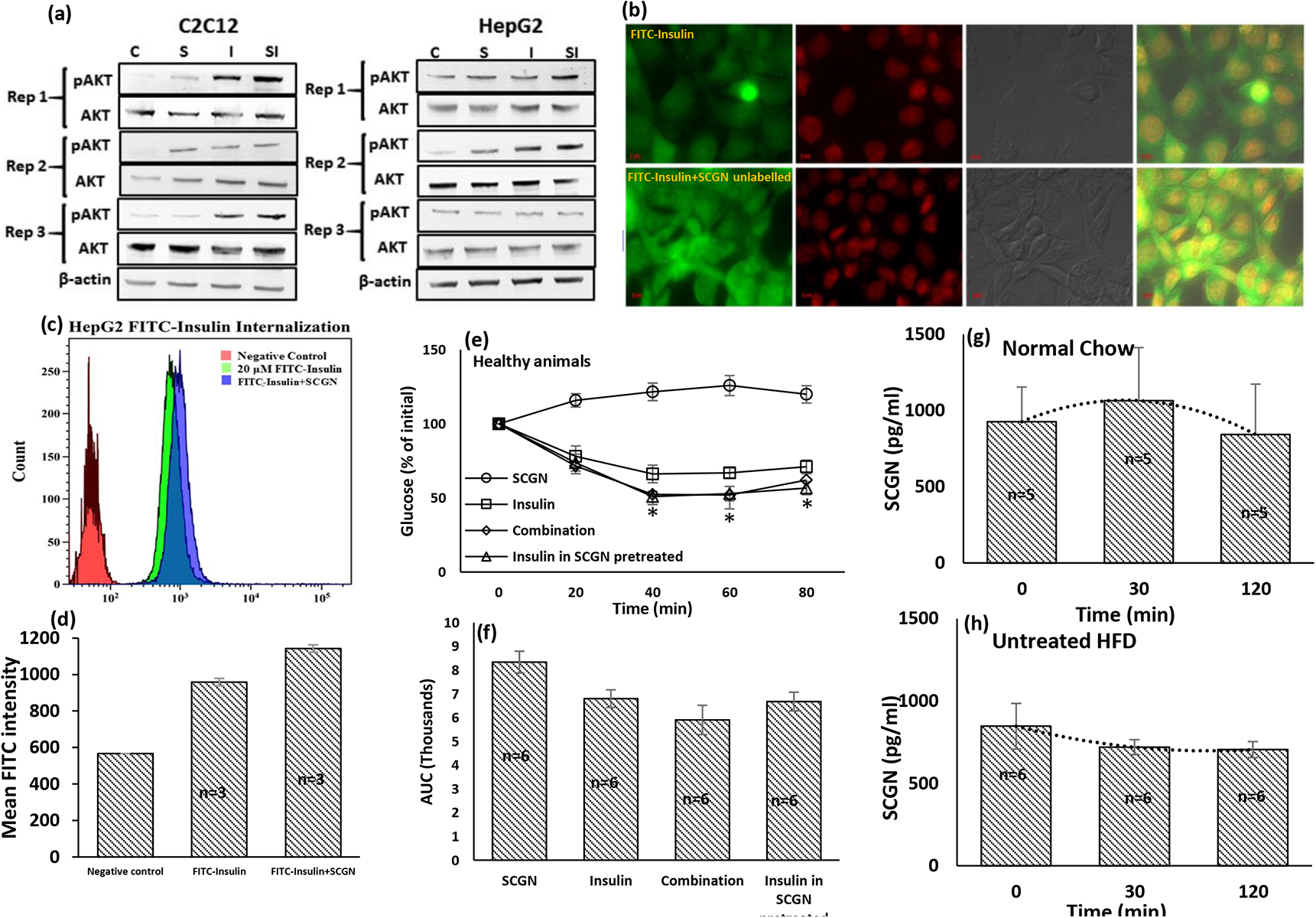
SCGN enhances insulin internalization and downstream signalling. (a) Western blots showing the increased insulin-induced phosphorylation of Akt in the presence of SCGN in C2C12 and HepG2 cells; (b) microscopic images showing increased internalization of FITC-insulin conjugate into HepG2 cells in the presence of unlabelled SCGN. FITC-labelled insulin (10 µM) and 10 µM FITC-insulin mixed with unlabelled SCGN were incubated for an hour followed by acid buffer wash, fixation by formaldehyde and subsequently mounted on slides. (c) FACS-mediated quantification of FITC-insulin conjugate internalization in the presence and absence of SCGN; (d) arithmetic mean of internalized FITC-insulin intensity; (e) ITT and (f) AUC of BALB/c mice injected with either Insulin, or combination of Insulin and SCGN after 6 hours fasting. Serum SCGN concentration of C57BL/6 mice kept on (g) normal chow; (h) high fat diet measured by ELISA at different time points in OGTT paradigm.

We then hypothesized that if these results are a consequence of biologically relevant and functional SCGN-insulin interaction, it should be reflected in increased insulin potency *in vivo*. To test our hypothesis, we performed an insulin tolerance test (ITT) in healthy animals. Each group of animals received one of the following treatments: (i) insulin, (ii) SCGN-insulin combination, or (iii) SCGN alone. We observed that SCGN-insulin combination was more effective in clearing blood glucose than insulin alone (Fig. 3e, f). Owing to the apparent lack of glucose clearance by SCGN alone, the enhanced potency of SCGN-insulin combination is attributed to SCGN-insulin interaction. Since ITT is a measure of the potency and sensitivity of insulin *in vivo*, results substantiate a possibility of the therapeutic potential of SCGN in non-insulin dependent diabetes (Type 2 diabetes mellitus).

Next, we tested if SCGN is synchronously secreted with insulin to augment insulin action. We observed that in the OGTT paradigm, serum SCGN fluctuates similar to insulin. The SCGN concentration rises at 30 minutes after oral glucose concentration while at 120 min, the increase in SCGN concentration subsides (Fig. 3g). In contrast, in the high-fat diet fed animals, this harmony was disturbed and no such correlation was observed (Fig. 3h). It suggests that in normal physiology, SCGN is co-secreted with insulin and deregulation of this synchrony affects insulin function as in diabetes.

### Chronic SCGN treatment preserves insulin and glucose tolerances and lowers hyperglycemia in an insulin-dependent manner

Insulin resistance is a physiological state where insulin, despite its presence, cannot induce the clearance of circulating glucose. To explore if, in the insulin-resistant state, the glucose lowering activity of insulin can be enhanced by supplementing with exogenous SCGN, we studied the effect of rSCGN administration in insulin-resistant animals. Since the diet-induced obese (DIO) animal model mimics the multifactorial human lifestyle associated diabetes than monogenic knockout animal models, we utilized DIO mice to examine the insulin-sensitizing activity of exogenous SCGN. We made an ab-initio chronic SCGN treatment model wherein the treatment group received a subcutaneous rSCGN injection every other day from the day1 of the exposure to the founding factor (i.e. HFD) while control groups (HFD and normal- chow) received PBS injection. After 12 weeks of exposure to HFD and concurrent treatment, we assessed the effect of SCGN treatment on insulin and glucose tolerance. We observed that upon exogenous insulin injection, rSCGN treated HFD animals cleared blood glucose more efficiently than control HFD despite the adversity of higher initial blood glucose level (t=0 in ITT) (Fig. 4a, b). To check if the same is true for endogenous insulin, we performed OGTT. Consistently, we observed better glucose clearance in SCGN-treated mice (Fig. 4c, d). Although the initial blood glucose of both treated and untreated HFD fed animals was high, however, SCGN treated animals efficiently cleared excess glucose from the blood and achieved a normoglycemia comparable to normal diet fed animals (Fig. 4c, d).

**Figure 4.**
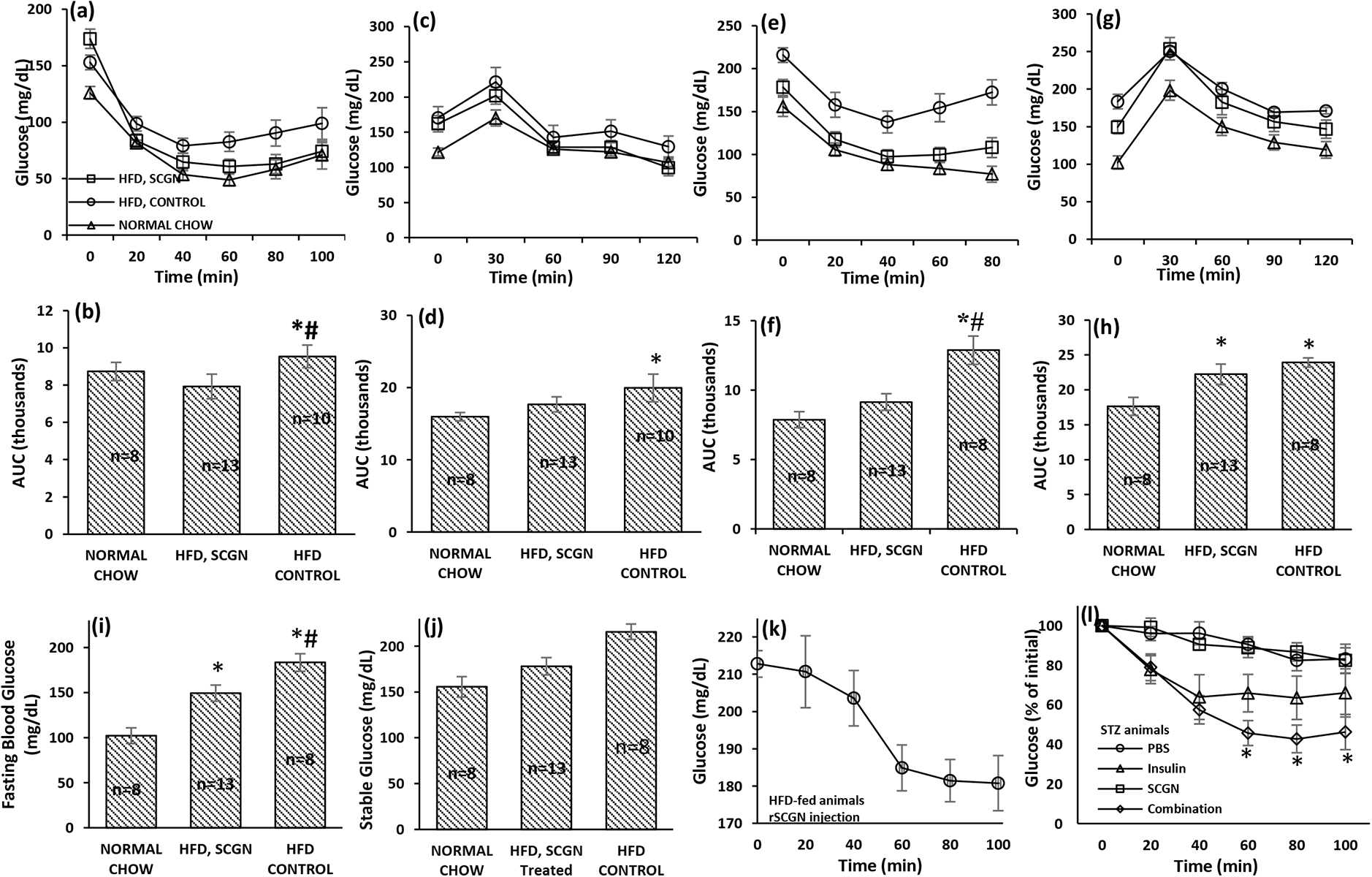
Chronic SCGN treatment alleviates insulin resistance in DIO animals. (a) ITT; (b) AUC of chow/HFD-fed animals after 12 weeks of exposure to HFD *P=0.002, #P=0.047; (c) OGTT; (d) AUC of the animals after 13 weeks *P=0.037. ITT and OGTT were repeated after additional two months of SCGN treatment (or vehicle injection). (e) ITT; (f) AUC, *P=0.0006, #P=0.004; (g) OGTT; (h) AUC, *P=0.0006 for chow vs untreated-hfd and *P=0.014 for chow vs SCGN-treated hfd. (i) Fasting blood glucose levels of DIO mice (0 min glucose values in last OGTT), *P=0.00001 for chow vs hfd control, *P=0.001 for chow vs hfd treated and #P=0.0115 for hfd treated vs untreated; (j) Stable blood glucose levels of DIO mice (0 min glucose values in last ITT), *P=0.0004 for chow vs untreated-hfd, *P=0.073 for chow vs treated-hfd and #P=0.006 for hfd-treated vs -untreated; (k) blood glucose after i.p. injection of rSCGN (10 mg/kg bw) in hfd-fed animals; (l) ITT in STZ animals after injection of SCGN or insulin or combination. *significance with respect to normal chow; # significance between treated and untreated hfd.

Encouraged by these results, we continued HFD (and concomitant rSCGN treatment) for additional two months to assess the efficacy of treatment till the severity of insulin resistance and hyperglycemia. At the conclusion of the experimental period, we found that SCGN-treated HFD animals had complete preservance of insulin sensitivity comparable to the chow-fed control animals demonstrating the anti-diabetic potential of rSCGN (Fig. 4e, f). Consistently, the glucose was better cleared by rSCGN treated HFD animals than untreated HFD group (Fig. 4g, h). The results are better reflected in stable blood glucose (derived from t=0 of ITT) and fasting blood glucose (derived from t=0 of OGTT) where the SCGN treated animals have lower blood glucose levels, reflecting a better glycemic control and sustenance of normoglycemia (Fig. 4i, j). In addition, rSCGN injection to HFD-fed animals induced an insulin dependent glucose clearance (Fig. 4k).

Secretagogin-induced lowering of glucose in HFD mice compelled us to explore if SCGN wields insulin mimic activity in vivo. We found that rSCGN injection to STZ animals had no significant glucose lowing effect in b-cell deficient STZ animals suggesting the lack of insulin mimetic activity of rSCGN (Fig. 4l). Nonetheless, SCGN-insulin combination was effective, (although a four weeks of alternate day rSCGN treatment was required to sensitize the animals) (Fig. 4l). However, in the mild hyperglycemic animals, pre-sensitization was not required. These results suggest that rSCGN does not have insulin mimic activity, rather rSCGN treatment increases insulin sensitivity and potency in both insulin-resistant and insulin-deficient animals.

### Chronic rSCGN injection relieves insulin resistance by alleviating hyperinsulinemia

A pronounced improvement in insulin tolerance of rSCGN-treated HFD-fed mice suggests better insulin sensitivity after SCGN treatment. Since the sustained hyperinsulinemia is both a cause and consequence of insulin resistance (27) and because SCGN is already shown to regulate expression and secretion of insulin (6, 16), we next checked if the reduced insulin level is one of the mechanisms by which SCGN alleviates insulin resistance. We found that the control chow group had the lowest insulin concentration while the HFD-fed untreated animals had highest circulating insulin at different time points in OGTT paradigm (Fig. 5a). The untreated HFD fed animals had highest circulating insulin concentration, suggesting the greatest degree of insulin resistance. In contrary, a better insulin tolerance and lower insulin concentration in rSCGN-treated HFD group supports higher insulin sensitivity (Fig. 5a). To calculate the extent of insulin sensitivity in control HFD group and to find the extent of protection exerted by rSCGN treatment, we calculated insulin sensitivity index, ISI_(0, 120)_ (28). We found that the normal chow fed animals had best insulin sensitivity; nonetheless, rSCGN treated mice had significantly higher ISI_(0, 120)_ suggesting better insulin sensitivity than untreated-HFD (Fig. 5b).

**Figure 5.**
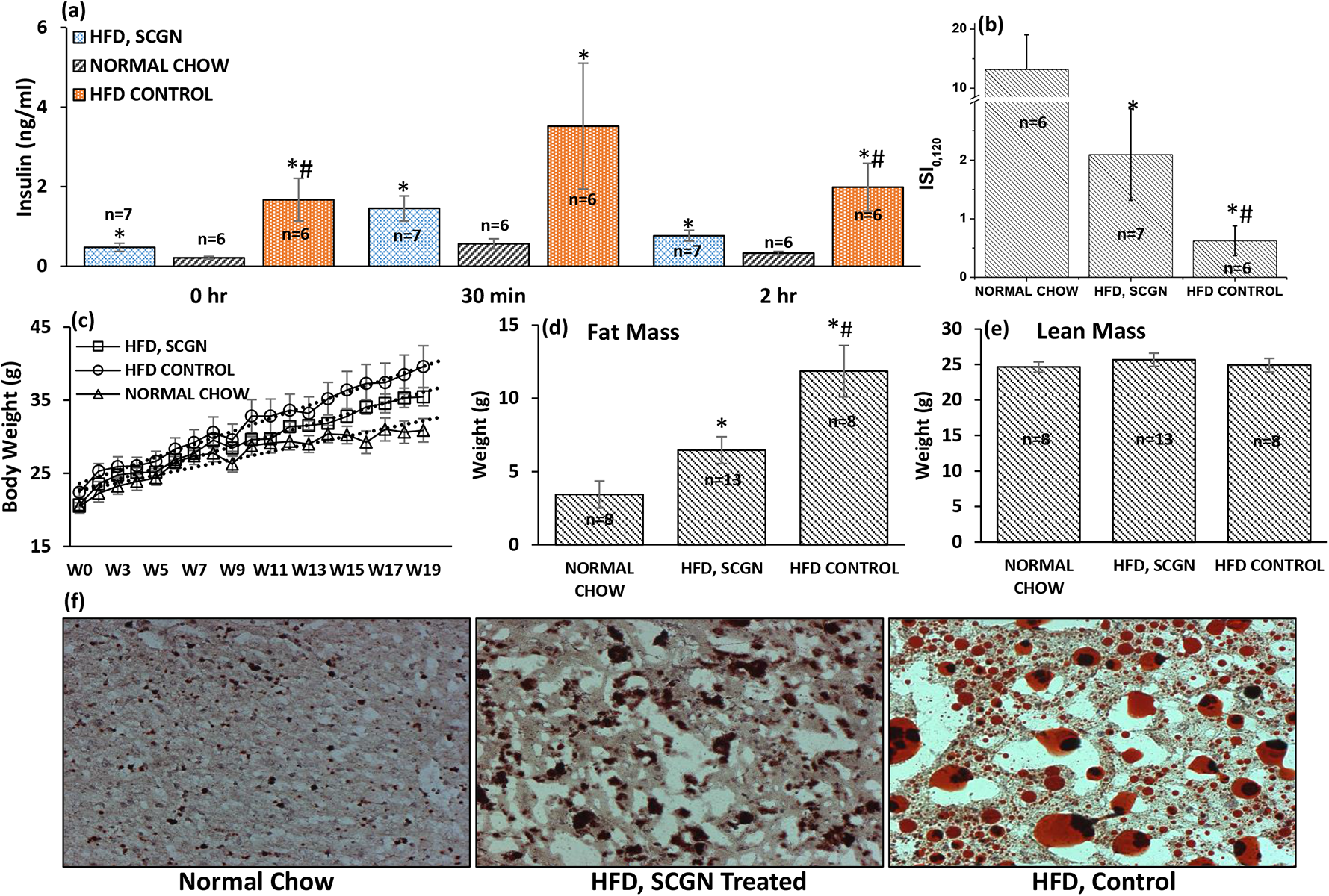
SCGN treatment moderates Hyperinsulinemia and fat deposition. (a) Serum Insulin concentration measured by ELISA in the same cohort of animals at different time points in OGTT paradigm, at 0 hr: *P=0.019 for chow vs hfd control, *P=0.022 for chow vs hfd treated and #P=0.018; at 30 min: *P=0.015; at 2 hours: *P=0.019 for chow vs hfd control, *P=0.007 for chow vs hfd treated and #P=0.027; (b) Insulin sensitivty index (ISI_0, 120_), *P=0.015 for chow vs untreated-hfd, *P=0.026 for chow vs treated-hfd and #P=0.048 for hfd-treated vs -untreated; (c) Change in body weight in the same cohort of animals over a period of 20 weeks; (d) Total body fat measured by echoMRI, *P=0.002 for chow vs untreated-hfd, *P=0.04 for chow vs treated-hfd and #P=0.011; (e) Lean mass (differences between the groups are insignificant); (f) Oil Red O staining of liver cryosections. *significance with respect to normal chow; # significance between treated and untreated hfd. All images were captured at 10x magnification and are presented with unaltered dimensions.

Obesity and increased body fat are major risk factors for lifestyle associated (such as diet) diabetes. Moreover, hyperinsulinemia also contributes to weight gain (29-35) which further increases insulin resistance. We, hence, followed the weight gain in animals on HFD. We observed that rSCGN treated animals had a lower rate of weight gain (Fig. 5c). In contrast, untreated HFD animals showed a higher degree of body weight gain (as apparent from the slop of incremental weight data). Consistently, the rSCGN-treated animals had significantly lower body fat than untreated HFD mice while the lean weight was comparable among groups as calculated from echo MRI measurements (Fig. 5d, e). In addition to reduced body fat, Oil red O staining suggested reduced hepatic triglycerides and lipid levels in rSCGN-treated animals than HFD control (Fig. 5f). The hepatic steatosis symptoms were alleviated in rSCGN treated animals with smaller lipid droplets and reduced necrotic area. These results suggest that rSCGN treatment alleviates insulin resistance likely by regulating body fat deposition and hepatic integrity.

### Systemic protection in DIO after SCGN treatment

Altered body fat composition leads to adjusted lipid metabolism, which precipitates the risk of cardiovascular diseases (36). We, thus, checked the effect of rSCGN treatment of serum lipid profile and other related factors (Fig. 6a-e). We observed that rSCGN treatment marginally reduced total serum triglyceride level while cholesterol levels were significantly reduced compared to untreated HFD-fed animals (Fig. 6a, b). The reduction in total cholesterol is likely due to reduction in LDL concentration because the HDL level remain comparable to untreated animals. (Fig. 6c-d) suggesting that rSCGN treatment might lessens the obesity-associated risk of cardiovascular diseases. Since SCGN is a Ca2+ sensor protein and because altered serum Ca2+ is a marker for early-stage insulin resistance (37-40); we also measured the serum Ca2+ concentration. We found that the rSCGN treated animals had lower serum Ca2+ reconfirming the insulin sensitization in rSCGN-treated HFD-fed animals (Fig. 6e).

**Figure 6.**
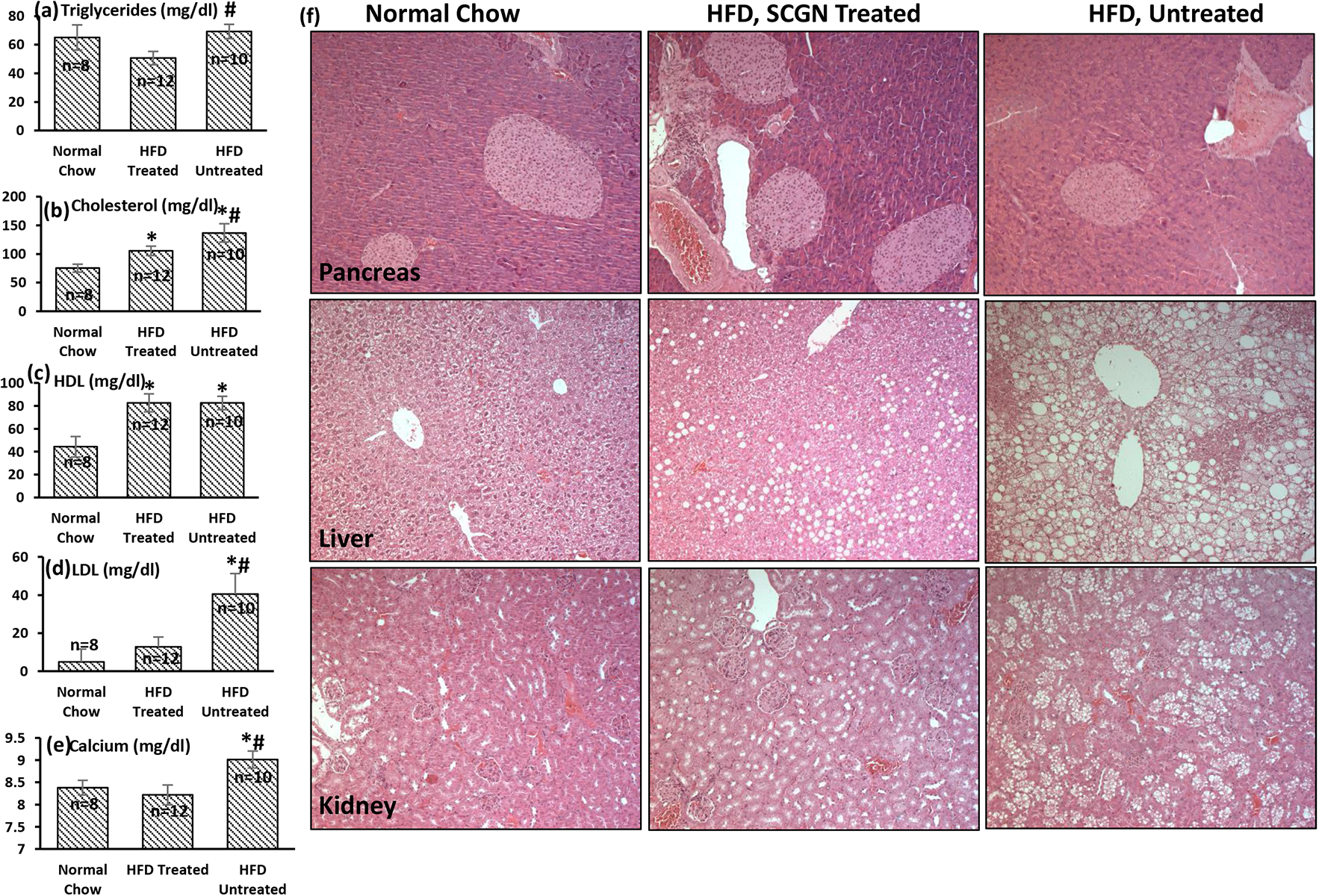
SCGN treatment shows no side-effects. (a) H&E staining of formalin fixed pancreas, liver and kidney sections. Serum analysis of (b) total triglycerides, #P=0.006; (c) cholesterol, *P=0.01 for chow vs untreated-hfd, *P=0.02 for chow vs treated-hfd, #P=0.04; (d) HDL-cholesterol, *P=0.001 for chow vs untreated-hfd, *P=0.003 for chow vs treated-hfd; (e) LDL-cholesterol, *P=0.024, #P=0.013; (f) Ca^2+^, *P=0.014, #P=0.007. All images were captured at 10x magnification and are presented with unaltered dimensions.

We next looked for tissue level modifications in the animals. We found that rSCGN treated animals had increased islet density and islet area than corresponding controls (fig. 6f). This observation extends the recent finding that SCGN promotes islet integrity and b-cell mass (16). The liver of untreated animals had prominent hepatic necrosis along with large fat bodies (as also visualized in Oil Red O stain) while treated animals were largely protected from necrosis and fat deposition. Similarly, kidney of untreated HFD-fed animals was severely degenerated seen as large necrotic area devoid of cells. The glomerular density was also reduced in untreated HFD-fed animals. In rSCGN treated animals, these features were less prominent likely due to reduced glycemic and metabolic load. These observations suggest that rSCGN treatment not only improves the glucose and lipid metabolism, it also helps keep metabolic organs healthy.

### SCGN treatment causes pancreatic b-cell regeneration and transcriptional reprogramming

Based on the our observation that rSCGN treatment increased the b-cell proliferation in HFD animals, we wondered if it will exert same effect in β-cell deficient STZ animals. We continued rSCGN injection (10 mg/kg, every alternate day) in STZ animals for four weeks. We observed that the control animals had expected less frequency of islets in pancreatic sections; however, rSCGN treated animals had increased islet frequency and the islet area (Fig. 7a). Although it is not clear if exogenous rSCGN injection mimics the endogenous molecular function to maintain the β-cell mass or embraces a new pathway, nonetheless, our results suggest a prospective application of SCGN as a regenerative medicine in T1D patients. While it is interesting to note a therapeutically relevant islet regenerative activity of rSCGN, it will be interesting to decern the active minimal peptide with enhanced efficacy and specificity at lower doses.

**Figure 7.**
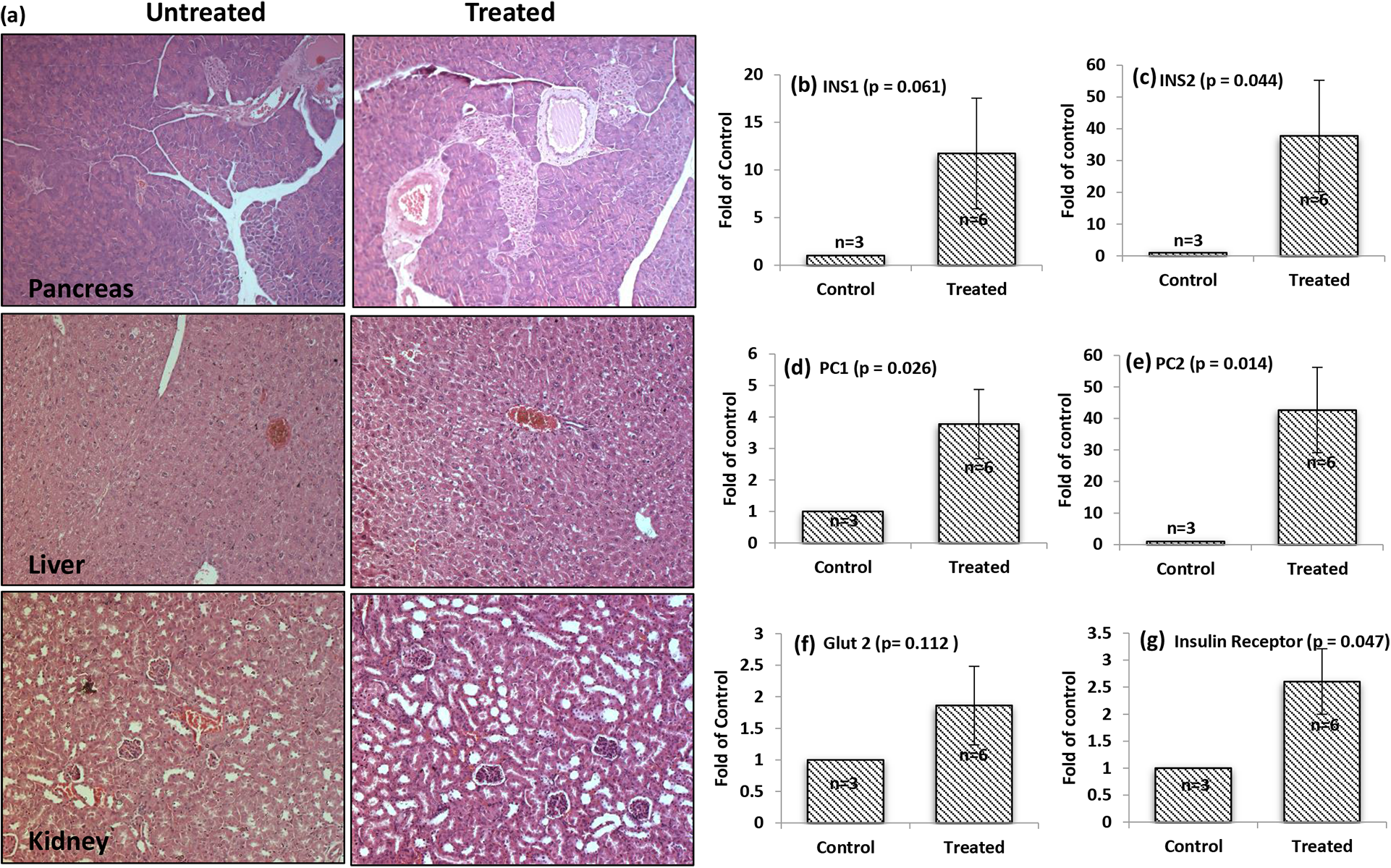
SCGN induced pancreatic β-cell proliferation and altered gene expression in STZ BALB/c mice. (a) H&E staining of formalin fixed pancreas, liver and kidney sections of STZ treated BALB/c mice upon SCGN treatment; (b-g) Transcriptional reprogramming of pancreatic genes in STZ untreated and SCGN treated animals as measured by qRT-PCR. All images were captured at 10x magnification and are presented with unaltered dimensions.

Beside pancreas, we observed that the SCGN treatment provides protection against kidney damage as well. The SCGN treated STZ animals, contrary to control STZ animals which had severe kidney damage with associated visible hemorrhagic-lesions, were generally protected from such damage and lesions (Fig. 7a). The liver of both treated and untreated groups was largely undamaged with infrequent macrophage invasion in the control animals (Fig. 7a).

To validate if the enhanced islets harbor functional β-cells, we characterized the molecular expression of β-cell specific genes by qRT-PCR analysis (Fig. 7b-h). We observed a substantial increase in the expression profile of insulin signaling related genes in pancreas, suggesting an increase in the total b-cell mass. Both the isoforms of insulin *(INS1* and *INS2)* were upregulated in the pancreas of SCGN-treated animals suggesting the true increase in the β-cell content (Fig. 7b, c). In addition, *INSR, GLUT2, PC1/3*, and *PC2* were significantly unregulated (Fig. 7d-h). Since Glut2 is the primary physiological sensor of glucose, increased expression suggests that the treated animals would have increased glucose responsive insulin release (41). The increased β-cell mass and enhanced insulin expression suggest that SCGN treatment can increase the insulin-producing capacity of insulin-deficient animals while the increase InsRec expression provides a way to achieve increased insulin sensitivity. PC1/3 is the critical enzyme for insulin maturation, upregulation of PC1 fulfills the obvious requirement of insulin maturation. Interestingly PC2, which is primarily involved in the glucagon maturation, was also upregulated in the SCGN treated animals. Though we did not explore the possibility, it suggests that SCGN may have a functional role in α-cell biology and glucagon secretion from the pancreas.

## Discussion

Insulin, a disulfide-linked two-chain hormone secreted from pancreatic β-cells, acts on target tissues (such as liver, muscles, fat tissue and brain) to maintain the normoglycemia. Because of its aggregation-prone β-chain and redox-sensitive disulfide bonds (42-43), it necessitates a structural stabilizer. In the pancreatic β-cells, Zn^2+^ stabilizes the tetrameric/hexameric form of insulin (44-46). Once the complex reaches circulation, Zn^2+^ dissociates from insulin to render it a biologically active monomer (47-50), which is more prone to aggregation and fibrillation and other anomalies. These glitches wreak fundamental science contradiction and pharmaceutical difficulties in insulin manufacture and insulin therapy (51-53). In this study, we establish that SCGN is an insulin-binding protein (InsBP) and discuss the role of SCGN-insulin interaction in insulin stability and function.

Protein fibrillation, including that of insulin, is a prominent anomaly associated with diabetes and multiple neurodegenerative diseases (20-24). SCGN mediated inhibition of insulin fibrillation suggests that in the extracellular SCGN could act as a stabilizer for circulatory insulin. Interestingly, SCGN has been reported to exert a protective effect against the Alzheimer’s disease (54-55). Since the Alzheimer’s disease is a fibril-associated disease, it inflicts the existence of SCGN’s anti-amyloidogenic effect to other relevant proteins. Therefore, SCGN seems to be a promising candidate against the amyloidosis associated diseases, such as neurodegeneration and Type 2 diabetes. Besides exerting protection from fibrillation, SCGN also modulates insulin internalization suggesting that SCGN could also regulate hepatic insulin clearance. However, a more comprehensive study of SCGN mediated insulin internalization needs to be performed to decipher the receptor-mediated and receptor-independent functions of insulin in the presence of SCGN. The SCGN mediated increase in the insulin’s half-life seems an unviable explanation because of the comparable time-trajectory of ITT in both insulin or SCGN-insulin combination groups. Nonetheless, a possibility of an insulin-sensitizing effect of SCGN in animals remains unchallenged. For now, the effect of SCGN administration seems to be a blend of both the tissue sensitization and insulin potentiation.

SCGN induced increase in insulin response in healthy animals prompted us to explore if the same will be true in diabetic animals as well. We observed that the very first dose of SCGN was effective in improving the insulin response, however, for the sustained insulin sensitization and glucose lowering, a chronic SCGN treatment is required. The insulin dependence of SCGN action was further validated in STZ induced model of insulin deficiency. In both the animal models, we found that a subcutaneous injection is more effective and sustained than intraperitoneal injection. We also studied the impact of SCGN on hepatic fat deposition and degree of insulin resistance to discover the mechanism of the sustained insulin sensitivity. Pre-established hyperinsulinemia causes the fat deposition and insulin resistance (29-35). We observed that SCGN treatment prevents hyperinsulinemia which conceivably regulates insulin sensitivity and a reduction in hepatic and body fat deposition. In addition to mentioned factors, transcriptional regulation of insulin-related genes’ expression might also contribute to SCGN mediated effects. The molecular mechanism of SCGN mediated impediment of hyperinsulinemia is likely attributed to the transcriptional regulation by SCGN followed by regulation of insulin secretion. In addition, hepatic insulin clearance could also contribute to a reduction in circulating insulin levels in rSCGN treated DIO animals. Considering previous studies that hyperinsulinemia is the crucial factor for the weight gain in DIO animals (33-35), reduced weight gain in rSCGN treated DIO animals could be attributed to the reduction in hyperinsulinemia. However, another possible line of future exploration could be the neuronal effect of SCGN in regulating satiety and energy expenditure. Moreover, the role of SCGN in regulating the release of other hormones (such as leptin and incretins) need to be established.

While looking at the histological integrity of pancreas, liver, and kidney, we observed a positive systemic influence on the key metabolic organs which are ultimately reflected in better glycemic control in SCGN treated animals. In the backdrop of a recent study reporting the role of intracellular SCGN in maintaining b-cell mass (16), it is promising to note that exogenous SCGN treatment has a positive effect on islet regeneration. Future studies are needed to explore and optimize the therapeutic application of SCGN induced b-cell regeneration. This study is the maiden exploration suggesting that SCGN could be used as a combination therapy with insulin in T1D to increase the insulin potency and also to stimulate of b-cell regeneration. Moreover, owing to its insulin sensitizing effect, SCGN becomes a potential candidate against T2D. Since SCGN is a relatively large protein, it will be interesting to identify individual domains responsible for a specific function such as b-cell regeneration or insulin sensitization.

Available anti-diabetic drugs have associated side-effects (56). Metformin, for instance, may cause severe gastrointestinal complication and its use is limited in renal insufficiency (57). Similarly, other insulin sensitizing drugs carry a possibility of weight gain, liver problem and cardiovascular risk (58). In this regard, SCGN seems to be safe with no apparent side-effects seen in rSCGN treated animals. In contrast, SCGN exerts beneficial effects such as reduced serum Ca2+, LDL, and hepatic steatosis and hence seems to reduce the risk of obesity-associated cardiovascular disease.

In summary, this study reports a novel role of SCGN in regulating insulin potency *in vivo* both in healthy as well as in diabetic animals. rSCGN treatment increases insulin sensitivity and lowers systemic diabetes associated complications. The rSCGN appears to be effective as monotherapy against insulin resistance while it can also be used in combination of insulin to increase the insulin efficacy while promoting the pancreatic regeneration over the course of treatment. While SCGN emerges as a potential anti-diabetic protein, future studies are required to analyze the therapeutic applicability.

## Material and Methods

### Antibodies

Anti-SCGN (bs11754R; Bioss), anti-insulin (bs0855R; Bioss), anti-phospho-Akt (Ser473) rabbit mAb (#9018S; Cell Signalling Technology), anti-Akt (pan) rabbit mAb (#4691S; Cell Signalling Technology), anti-β-actin (bs0061R; Bioss), secondary HRP (bs0295G; Bioss), Alexa Fluor 594 Goat anti rabbit IgG (A11037; Life technologies Molecular probes).

### Pull down assay

His-tagged SCGN (100 μg) was allowed to bind to a Ni-NTA resin column in the presence of Ca^2+^. MIN6 cells were scrapped in RIPA buffer and sonicated for 10 cycles of 10 sec on/off cycle on a Sonics VibraCell sonicator at 20% efficiency. The whole MIN6 cell lysate was loaded on an SCGN-bound Ni-NTA resin column, washed with PBST buffer and eluted with 250 mM imidazole. The supernatant, wash, and eluents were resolved on 12% SDS-PAGE gels and gel pieces containing protein bands were excised. The samples were processed for mass spectrometry identification by desalting with zip-tip and analyzed on Q-Exactive Thermo Scientific. Data analysis was performed using Xcalibur software with a high stringency.

### Cell Culture

MIN6 cells were cultured in DMEM supplemented with 10% FBS, penicillin, and streptomycin. HepG2 cells were cultured in similar conditions. C2C12 myocytes were maintained in 20% serum and antibiotics. To differentiate, C2C12 cells were grown on a serum free DMEM media containing 100 nM insulin. After 24 hours, cells were grown in serum/insulin-free DMEM. The differentiation was assessed by tubular morphology and multinuclear structure by DAPI staining.

### Colocalization

SCGN was cloned in a pEGFP-N3 vector with C-terminal eGFP in-frame by restriction-free (RF) cloning. MIN6 cells (0.2x10^6^) were seeded on coverslips in a 6-well plate. Next day, transfection was done using Lipofectamine LTX Plus according to manufacturer’s protocol. After 24 hours, cells were visualized for the SCGN-eGFP expression and were fixed in 4% formaldehyde. Subsequently, the cells were permeabilized followed by blocking with 2% BSA at room temperature for 1 hour. Next, the cells were incubated with the anti-insulin antibody (1:500) for one hour, washed thrice with PBST (0.1% Tween 20) and incubated with Alexa Fluor

594 conjugated secondary antibody (1:500) for 1 hour followed by 5 washes with PBST. DAPI was used as a nuclear staining dye. Images captured on an Apotome 2 microscope, were adjusted for background and contrast using Image J software without manipulating any specific feature or part of the image.

### CoIP and *in vitro* binding assay

MIN6 cells were grown in a T75 tissue culture flask and trypsinized upon confluency. The cells collected after slow centrifugation were resuspended in lysis buffer (PBS, 0.1% Tween 20, protCEASE 50; G-Biosciences), and were ruptured by three freeze-thaw cycles and supernatant was used for immunoprecipitation. For IP from media, 10 ml of conditioned media (from a T75 flask in which 80-90% confluent MIN6 cells were cultured for 24 hours) was concentrated using Amicon Ultrafiltration device (cut-off of 3 kDa). Subsequently, five micrograms of anti-insulin antibody (or 5 µg of irrelevant IgG) was added to 1 ml cell lysate (or media) and incubated overnight at 4 ^°^C with continuous rotation. Antibody complex was precipitated using Protein A Sepharose beads (Amersham Biosciences) followed by 5 washes with PBST. The resin was boiled for 2 minutes in SDS-PAGE loading buffer centrifuged and the supernatant was resolved on 12% SDS-PAGE. The proteins were transferred to PVDF membrane and developed with the anti-SCGN antibody (1:10,000).

### Recombinant protein purification

Recombinant SCGN was purified from soluble fractions by using Ni-NTA column followed by gel filtration by modifying the earlier protocol (59). Briefly, SCGN expressing vector carrying E. *coli* cells were grown in minimal media. After 10 hours of post induction () incubation at 25 C, cell pellet was lysed in Buffer A (50 mM Tris, pH 7.5 and 100 mM KCl). After sonication and centrifugal clearance, supernatant was loaded onto a Ni-NTA column. Bound protein was washed with 8 column volumes of wash buffer 1 (50 mM Tris, pH 7.5, 150 mM KCl) then with 10 column volume of wash buffer 2 (50 mM Tris pH 7.5, 150 mM KCl, 2% Triton X100) followed by wash with wash buffer 1. Protein was eluted with a gradient flow of elution buffer (50 mM Tris, 150 mM KCl, 50-250 mM imidazole gradient). For preparing the Ca^2+^ free protein, gel filtered protein solution was incubated with 10xmolar concentration of EDTA for half an hour followed by buffer exchange with Chelex-purified buffer (50 mM Tris, pH 7.5, 150 mM KCl). For animals and cell culture, protein buffer was PBS.

### Protein overlay assay

Proteins were immobilized on a PVDF membrane with a slot-blot manifold apparatus (Amersham Biosciences). After blocking with 5% BSA, the membrane was incubated with 10 µM SCGN for one hour and washed four times with PBST. The further procedure was followed as standard Western Blotting with the anti-SCGN antibody (1:10,000).

### Bio-layer interferometry

BLI experiments were performed on an Octet Red 96 system. Our efforts to immobilize His-tagged-SCGN using Ni-NTA chemistry or using amine-coupling chemistry were unsuccessful because of the non-specific binding of insulin to either probe. We hence immobilized insulin (100 µg) on the probe using amine-coupling and 200 µl SCGN (100 µM) was taken as ligand in the wells in the absence or presence of 1 mM Ca^2+^. Initial baseline- 60 s, association/dissociation- 300 s each followed by a regeneration step. Two controls were run (i) insulin-immobilized probe and buffer in the well, (ii) free inert probe with SCGN in the well. There was an inappreciable signal in the control (i). Reference (ii) was subtracted from the sample. The data fitting was performed for the initial 120 seconds of association phase and 300 seconds of dissociation phase in the vendor-supplied module using 1:1 model. The R^2^ of >0.95 and Chi^2^ of <1 was used as the criteria of a good fit.

### Spectral measurements

Circular dichroism (CD) spectra were recorded on a Jasco J-815 spectropolarimeter, using 1 cm path length cuvettes. Fluorescence emission spectra were recorded on an F-4500 Hitachi spectrophotometer at an excitation wavelength of 295 nm. The excitation and emission band passes were set at 5 nm and spectra were recorded in 50 mM Tris (pH 7.5), 100 mM KCl. The spectra were corrected for insulin contribution. Near-UV CD spectra were recorded using 1 mg/ml protein concentration, and fluorescence spectra were recorded using 0.1 mg/ml protein (unless otherwise stated in the graph).

### Analytical gel filtration

Differential elution of free proteins and the complex was studied on a Superdex 75 (10/300) column (Wipro GE Healthcare) under described conditions. Column was pre-equilibrated with buffer (50 mM Tris, pH 7.5, 100 mM KCl) containing either 0.1 mM EDTA or 2 mM Ca2+.

### Insulin amyloidogenesis by TEM

Either the proteins or the complex were incubated overnight at 60 ºC without stirring. Fibrils thus formed were mixed. The solution was applied onto a 300-mesh copper grid and 2% freshly prepared uranyl acetate was added for staining. Dried samples were observed under a transmission electron microscope (Hitachi, Japan) operating at an accelerating voltage of 200 kV.

### Insulin amyloidogenesis and aggregation by fluorescence spectroscopy

For the real-time insulin fibrillation measurement, 0.5 mg/ml insulin (50 mM glycine, 100 mM KCl, pH 2.5, 10 µM ThT) was incubated at 55 ºC (in the absence of Ca^2+^) or 60 ºC (in the presence of Ca^2+^) with continuous stirring. ThT fluorescence was monitored for one hour (λ_ex_ =440 nm, λ_em_=482 nm). For DTT-induced aggregation, 0.2 mg/ml insulin (in 50 mM Tris buffer pH 7.5, 100 mM KCl) in the presence or the absence of different concentrations of SCGN was equilibrated at 37 °C for 10 min with constant stirring in the cuvette. The reduction of insulin was initiated by the addition of 20 mM DTT to the sample. The extent of aggregation was monitored as scattering (excitation and emission monochromators set at 465 nm) set at 5 nm band passes in time scan mode of the instrument.

### MTT assay

Fibrils were generated by overnight incubation of 1 mg/ml insulin in 50 mM glycine pH 2.5, 100 mM KCl buffer at 65 ºC. MIN6 cells were seeded at a density of 20,000 cells/well in a 96-well plate. Next day, the cells were washed with PBS and incubated with 10 µM SCGN, 20 µM insulin fibrils (fibrils), complex of 10 µM SCGN and 20 µM fibrils, 5% Triton X-100 (negative control), or BSA and fibrils complex in serum-free media for 24 hours. Cells were washed again with PBS and incubated with 100 μl of 5 mg/ml MTT in incomplete media for 4 hours at 37 ºC. Fifty microliters of media were replaced with DMSO and mixed properly to dissolve crystals. The absorbance was recorded at 540 nm on a Perkin-Elmer multimode Plate reader. The number of samples per treatment was eight (n=8).

### FITC-Insulin preparation

Five-milligram insulin powder was dissolved in 1 ml glycine buffer (50 mM glycine, 100 mM KCl, pH 2.5, 1 mM EDTA) to avoid insulin aggregation. After insulin was dissolved, the solution was diluted 4 times. In parallel, 2 mg FITC was dissolved in 300 µl ethanol. FITC was added to the insulin solution dropwise on a magnetic stirrer in dark at room temperature. After 3 hours of the coupling reaction, the solution was subjected to a size exclusion column attached to a BioRad FPLC system. Pure insulin fractions were decided by the absorbance at 280 and 495 nm set in the Quadtec detector. The extent of FITC conjugation was calculated exactly as recommended by the SIGMA (CAS #: 3326-32-7). Our preparation was found to have ~4 FITC molecules per insulin molecule.

### Microscopic and FACS study of FITC-insulin internalization

HepG2 cells were seeded on the coverslips in a six-well plate (0.2x10^6^ cells/well). After cells reached ~80-90% confluency, the cells were serum deprived for 24 hours and were treated with 10 µM FITC-insulin either in the presence or absence of 10 µM of unlabelled SCGN. After 30 min incubation, the culture plate was transferred to the ice and was washed thrice with ice-cold PBS followed by two washes with ice-cold glycine-NaCl buffer (120 mM glycine, 150 mM NaCl, pH 2.5) to remove the membrane-bound FITC-insulin. After additional two washes with ice-cold PBS, cells on coverslips were fixed and mounted on a glass slide with DAPI containing mounting media. For the better visualization, DAPI was pseudo-colored to red instead of conventional blue color. The images were captured on an Apotome 2 system with 63x objective and oil immersion medium. The exposure time for each channel was constant for all the samples. No image processing was performed on the acquired images.

For FACS-assisted quantification of FITC-insulin internalization, samples were prepared similarly except that coverslips were not used for FACS samples. After the washes, the cells were trypsinized, centrifuged and the cell pellet was resuspended in 400 µl PBS. Immediately, FACS analysis was performed on a Beckman Coulter Gallios Flow Cytometer system using FITC channel. At least 20,000 cells were analyzed for each sample. The data analysis and figure preparation were performed on the Kaluza software module provided by the vendor.

### Akt phosphorylation assay

C2C12 cells were differentiated in T25 cell culture flasks. After glucose deprivation (2 hrs for C2C12 and 4 hrs for HepG2 cells) in glucose-free KRP (20 mM HEPES, 0.4 mM K_2_HPO_4_, 1 mM MgSO_4_, 5 mM KCl, 135 mM NaCl), the cells were washed thrice and treated with insulin (100 nM) or SCGN (100 nM) or the complex of the two in KRP buffer. Control samples received the corresponding buffer. After 20 minutes, cells were lysed in RIPA buffer. Total protein was quantified by BCA method and 50 µg protein was resolved on 12% SDS-PAGE followed by Western blotting. First, pAkt was probed against Akt phosphorylated (Ser473) antibody (1:10,000) followed by stripping of the membrane and co-detection of β-actin (1:10,000) and Akt (1:10,000) with respective antibodies.

### Animals maintenance, OGTT and ITT

Animals (after due approval from Institutional Animal Ethics Committee, 91/2015, 2/2017 and 35/2017,) were kept in Individually Ventilated Cages on normal chow (*BALB/c* mice) or on high fat diet (*C57BL/6J* mice). To induce diabetes in *BALB/c* mice, STZ (100 mg/kg bodyweight in citrate buffer pH 4.5) was injected intraperitoneally after six hours of fasting. After 3 days, glucose was measured and animals showing random glucose >200 mg/dL were selected for experiments. *C57BL/6J* mice were kept on high fat diet (HFD; D12492, Research Diets, New Brunswick, NJ) or normal chow and the rSCGN treatment was started simultaneously. After three months, first set of OGTT and ITT experiments were performed.

OGTT was conducted after overnight fasting. Mice were given an oral bolus of 2 g glucose per kilogram body weight and blood glucose was monitored at every 30 min using a AccuCheck active OneTouch glucometer (Roche).

ITT was conducted after 6 h of fasting. Mice were injected with (i.p.) 0.75 IU of fast acting insulin per kilogram body weight and blood glucose was monitored at every 20 min using a AccuCheck active OneTouch glucometer (Roche).

### Serum Analysis

Random triglyceride (GPO/PAP method, TG kit, Coral Clinical Systems), Cholesterol (CHOD/PAP method, CHOL Kit, Coral Clinical Systems), HDL (Direct enzymatic method, HDL-D kit, Coral Clinical Systems), Ca^2+^ (OCPC method, CAL Kit, Coral Clinical Systems) and Glucose (Glucose (GO) assay kit; cat: GAGO20-1KT, SIGMA) measurements were performed as per the product guidelines and protocols. LDL was calculated by the equation: (Total Cholesterol) - (HDL) - (TG/5).

### ELISA

Insulin (Ultra-Sensitive Mouse Insulin ELISA kit, Crystal Chem; cat: 90080) and SCGN (Secretagogin BioAssay^TM^, USBiological; cat: 027968) measurements were performed as per the vendor’s guidelines following the product protocols.

### Insulin sensitivity index [ISI_(0,120)_]

The insulin sensitivity index was calculated precisely as described (28). Briefly, after 16 hours of fasting, blood was drawn for serum insulin/glucose measurement. Two hours after the oral glucose bolus (as described in OGTT), another sample of blood was collected. Serum glucose and insulin levels were determined as described in respective sections. MCR and MCI were calculated and insulin sensitivity was calculated as ISI_0,120_= MCR/log MSI.

### Oil Red O staining

Frozen liver tissues were sectioned on a microtome at the thickness of 5 micrometers and immobilized on charged slides. Oil Red O (SIGMA, O1391-250ML) staining was performed as follows. Tissue were wet in distilled H_2_O followed by a wash with PBS. Then slides were dipped in 60% isopropanol followed by immersion in Oil RedO solution for 1 hour. Next, slides were washed twice with PBS (5 min each) followed by immersion in hematoxylin solution for 4 minutes. Finally, slides were washed in running tap water and glycerol soaked tissues was covered with coverslips. Slides were observed under Axioplan 2 imaging system.

### Echo-MRI

EchoMRI™-500, which is designed to analyze the body composition for live small animals, was used for total fat content measurement. Animals were kept in vendor provided animals restrainer. All readings were the average of three accumulations.

### Histology

At the end of experiments (and after a recovery period), animals were sacrificed using recommended procedures. Tissues were removed and fixed in formalin (10% formaldehyde in phosphate-buffered) followed by embedding. The paraffin embedded tissue were sectioned at 10 micrometer thickness. Sections were stained with hematoxylin and eosin and were visualized on Axioplan 2 imaging system.

### qRT-PCR

Tissue samples, immediately after removal from animal, were washed and immersed in TRIzol RNAiso plus reagent and the tissue was stored at -80 ºC. At the time of processing, tissues were thawed and total RNA was isolated by chloroform extraction. Upon checking the integrity and concentration of RNA, 5 µg of RNA was used for cDNA preparation using SuperScript III First-Strand Synthesis System (Thermo Fisher Scientific). mRNA levels were quantified by qRT-PCR (ABI Prism 7900 HT; Applied Biosystems) using SYBR Green (Applied Biosystems). Primer sequences are available on request.

### Statistical analysis

All significance test for group-wise comparison were performed on MS excel using student’s t-test except OGTT and ITT time course measurement where two-way anova analysis (with Holm-Sidak post-hoc analysis) was performed on sigma plot software.

## Acknowledgements

We are grateful to Dr. B Raman for help in Biolayer interferometry and Dr. Veena Yeruva for help in initial blotting experiments and occasional chemical. Mr Prashant has been helpful in few animal experiments. Mr. Harikrishna helped in TEM imaging. We acknowledge the Central Tissue Culture, Fine Bio-Chemicals, Microscopy, Animal House, and Mass Spectrometry facility of CCMB. Syed Sayeed Abdul for excellent laboratory assistance and Dr. Sushil Chandani for discussion and critical reading of the manuscript. In addition, discussions with Dr. GR Chandak and Dr. Jyotsna Dhawan were useful in experimental design.

## Author Contributions

Inception: AKS; Experimental design: AKS, RK, JMK, YS; Data acquisition: AKS, RK, SC, SR, AR; Data analysis: AKS, RK, JMK, YS; Resources: YS; Writing - original draft: AKS, RK, YS; Writing - review & editing: AKS, RK, JMK, YS; Supervision: YS; Project administration: YS; Funding acquisition: YS.

